# Developmental Alterations in Brain Network Asymmetry in 3- to 9-Month Infants with Congenital Sensorineural Hearing Loss

**DOI:** 10.1101/2023.05.31.543147

**Authors:** Guangfang Liu, Xin Zhou, Zhenyan Hu, Yidi Liu, Endi Huo, Heather Bortfeld, Qi Dong, Haihong Liu, Haijing Niu

## Abstract

Auditory exposure plays crucial roles in shaping healthy brain development and generating lateralization of functional network organization. However, little is known about whether and how an initial lack of auditory exposure in early infancy may disrupt development of functional network lateralization. We addressed this issue by recruiting 55 infants with congenital sensorineural hearing loss (SNHL) and 60 typically developing (TD) controls. Resting-state fNIRS imaging data were acquired to construct hemispheric cerebral networks, and graph theory was applied to quantify the topological characteristics of hemispheric networks. The infants with SNHL exhibited efficient small-world characteristic within each hemispheric network, however, the lateralization of functional network efficiency was substantially disrupted. Compared with TD infants with significantly increased network efficiency lateralized toward left hemisphere with age, the SNHL infants did not exhibit the emergence and development of such cerebral lateralization. Furthermore, the increased leftward asymmetry in nodal efficiency with age was found in TD but not in SNHL infants. Interestingly, the degree of hearing loss had no correlation with lateralization strength in the SNHL group. These results suggest that SNHL infants exhibited disrupted development of cortical lateralization in functional network organization, and highlight the importance of auditory stimulation-promoted multisensory functional integration in early infancy.

## 1 Introduction

Sensorineural hearing loss (SNHL) is the most prevalent congenital sensory deficit in neonates. Approximately 0.1% ∼ 0.3% of newborns suffer from complete or partial loss of hearing at birth in Western countries (Kral and O’Donoghue 2010; Ospina-García et al. 2019). Deprivation of early auditory experiences can profoundly influence infants’ language development, executive function, personality, and social behavior (Bavelier et al. 2000; Figueras et al. 2008; Tomblin et al. 2015).

A growing body of neuroimaging evidence demonstrates that the brains of infants with SNHL exhibit altered structural and functional connectivity in multiple brain regions. For instance, a study by Wang et al. (2019) revealed widespread white matter alterations in the auditory and language regions of the brain (for example, bilateral superior temporal gyrus) and increased functional connectivity within these regions. Another study reported that SNHL infants showed enhanced functional connectivity within the auditory and salience networks, while functional connectivity within the default mode network was decreased (Wang et al. 2021). Additionally, Xia et al. (2017) found functions in the hearing and language regions were impaired and function in the vision regions was enhanced in the SNHL infants. These findings indicate that the loss and the compensatory reorganization of brain network functional connectivity coexist in SNHL infants. However, few studies to date have focused on how the brain functions of the left and right hemispheres develop in SNHL infants.

Hemispheric asymmetry refers to how the two hemispheres operate in different, albeit integrated ways; it is considered one of the cardinal features of human brain development (Hutsler & Galuske, 2003; G., Toga & Thompson, 2003), and age-associated increases in functional integration efficiency are observed across early development. Meanwhile, evidence has also revealed that hemispheric asymmetry is disrupted in individuals with SNHL (Liu et al. 2022). Marcotte and Morere (1990) found that adolescents with congenitally deafness or who acquired deafness early in life (onset between 6 and 36 months of age) showed anomalous cerebral representation, especially atypical lateralization. Twomey et al. (2017) found that participants who were deaf from birth show atypical brain lateralization when they process auditory stimuli. Of note, participants in these studies represented wide age ranges, and the overall cognitive function and the influence of rehabilitation training are notable confounds. Because infancy is a critical period when the brain undergoes crucial changes from complex interactions between the internal and external environments, it is important to explore the impact of early hearing experience loss on the emergence of hemispheric asymmetry itself. Regrettably, it remains unknown how hemispheric network asymmetry develop in infancy in individuals with congenital SNHL.

Here, we investigated the developmental changes in topological asymmetry between the two hemispheres in congenital SNHL infants between 3 and 9 months of age, and compared these changes to those in age-matched TD infants. We hypothesized that SNHL infants would show atypical development of functional topological asymmetry of the cortical network. To test this hypothesis, we collected resting-state fNIRS data from 105 infants (55 SNHL infants and 60 TD infants). For each infant, hemispheric functional networks were constructed, and then topological network efficiency metrics were calculated utilizing graph theoretical approaches. Finally, the developmental trajectories of topological asymmetry between groups were characterized. To our knowledge, this is the first study to reveal developmental changes in hemispheric network asymmetry in congenital SNHL in early infancy. The findings will serve as a foundation for a better understanding of cortical development across two hemispheres in congenital SNHL infants.

## 2 Results

### 2.1 Efficient small-world properties of the hemispheric networks for SNHL and TD infants

For each hemisphere, the group-averaged functional connectivity and small-world attributes are shown in Figure 1 for SNHL and TD infants, respectively. Visually, the infants with SNHL showed widespread decreased functional connectivities within each hemisphere compared with TD infants (Figure 1A); however, the brain network organization in SNHL infants, similar to the TD infants, exhibited typical small-worldness properties. Specifically, the network global efficiency derived from the brain networks was approximately equal to that of their matched random networks for both SNHL and TD infants, and the network local efficiency derived from the brain networks was larger than that of their matched random networks. These patterns led to the normalized global efficiency being approximately equal to 1 and the normalized local efficiency being greater than 1, typical features of small-world topology. The small-world organization indicates that brains in both SNHL and TD infants were efficient in distributed information integration and local information processing.

**Figure 1.**
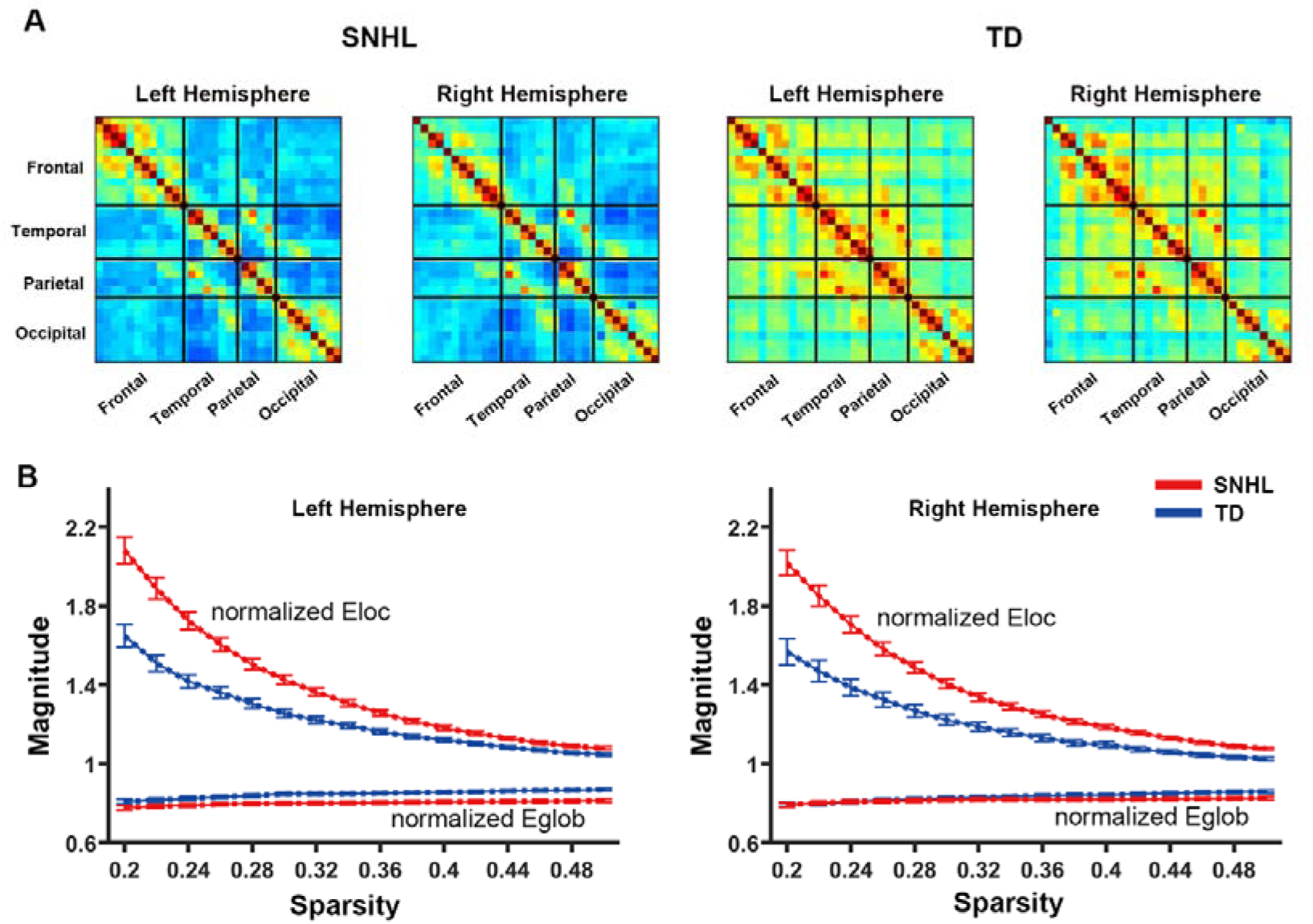
Functional connectivity maps and small-world network characteristics of the functional networks in TD and congenital SNHL infants. (A) Functional connectivity maps. (B) Small-world properties of the functional networks. Error bars indicate the standard errors of each subgroup.

### 2.2 Hemispheric asymmetry of global and local network efficiencies in each group

For each group, we examined the within-group hemispheric asymmetry after controlling the age factor. The TD infants showed significant leftward (i.e., left > right) hemispheric asymmetry in both global efficiency [t = 2.82, p = 0.007] (Figure 2A) and local efficiency [t = 2.13, p = 0.038] (Figure 2B). However, for infants with congenital SNHL, neither global nor local network efficiencies showed significant differences across the two hemispheres [*E_glob_*: t = 0.57, p = 0.57; *E_loc_*: t = 0.85, p = 0.40] (Figure 2). These results indicate that the left hemispheric dominance of the functional network organization was significantly disrupted in infants with congenital SNHL.

**Figure 2.**
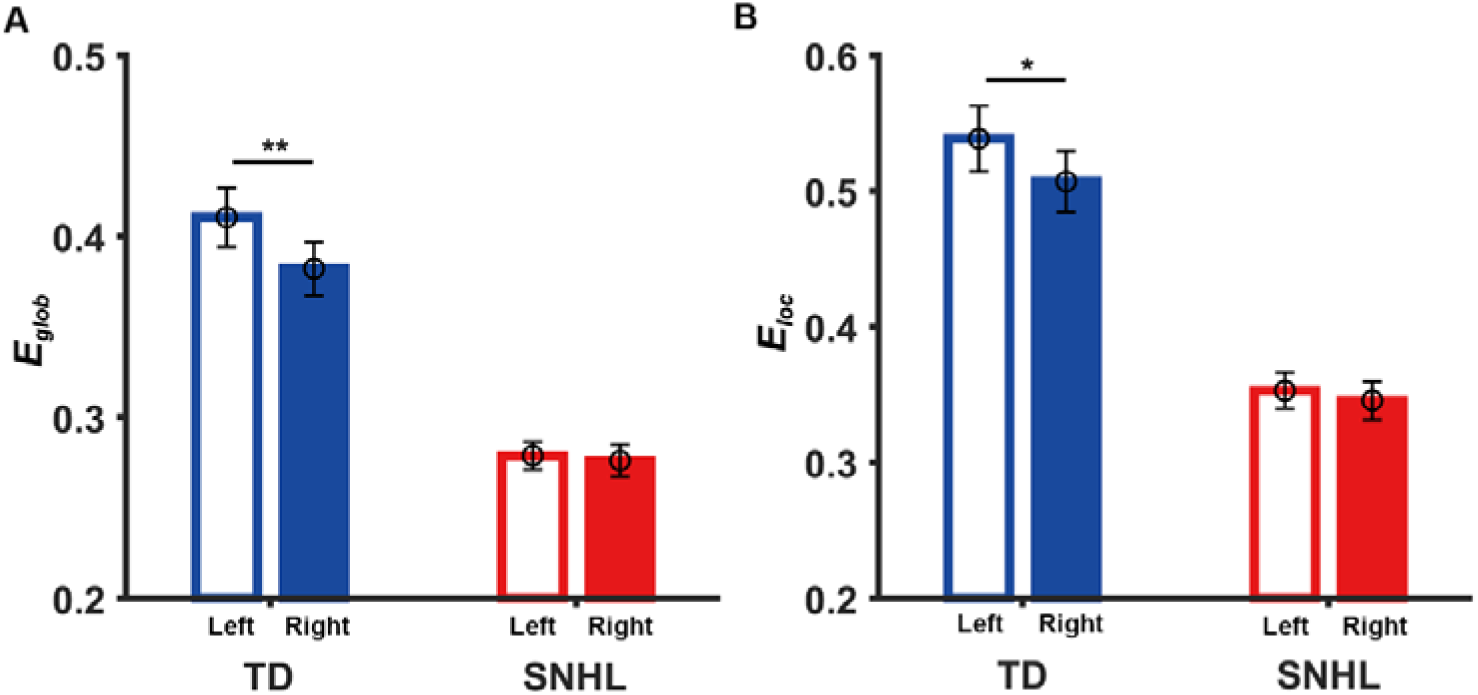
Within-group asymmetry in global efficiency (*E_glob_*) (A) and local efficiency (*E_loc_*) (B) for the TD infants and the SNHL infants. *p < 0.05, **p < 0.01.

### 2.3 Developmental differences of hemispheric asymmetry in global and local efficiencies across groups

We further compared the developmental difference of cortical network asymmetry in infants with congenital SNHL to that in TD infants based on the *AI* of efficiency measures. Statistical analysis from a general linear model revealed that there were significant group × age interaction effects on the *AI* of global efficiency [t = 2.38, p = 0.027] but not on local efficiency [t = 1.70, p = 0.091]. Post hoc analysis revealed that the TD infants showed a significant developmental increase in the asymmetry of global efficiency [r = 0.31, p = 0.015] and a trend toward an increase in the asymmetry of local efficiency [r = 0.21, p = 0.099]. Nevertheless, the infants with congenital SNHL did not show any developmental changes in the hemispheric asymmetry of either global or local network efficiency [*AI* of *E_glob_*: r = -0.05, p = 0.69; *AI* of *E_loc_*: r = -0.09, p = 0.52] (Figure 3A).

**Figure 3.**
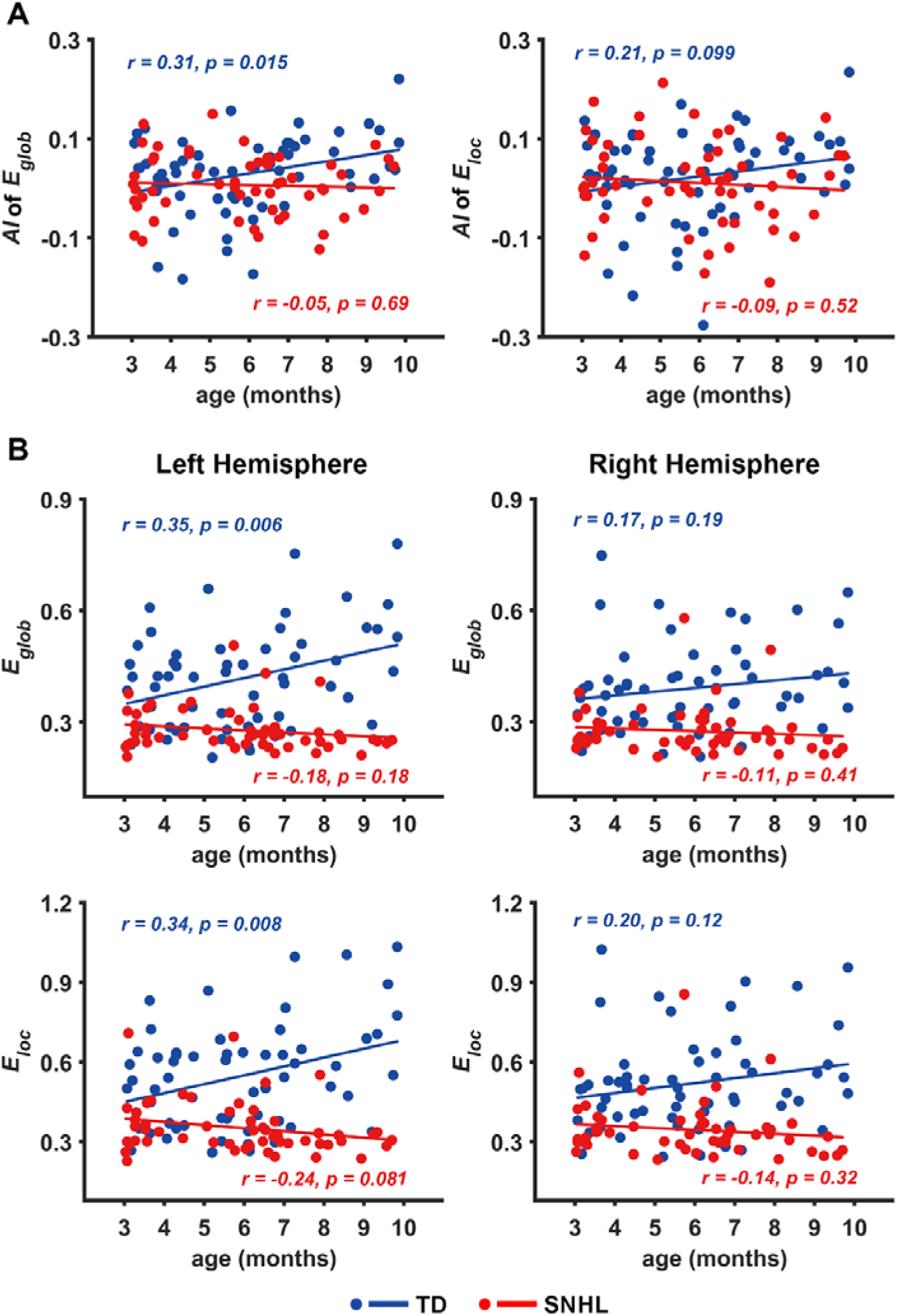
Developmental differences in hemispheric networks in TD and congenital SNHL infants. (A) Developmental comparisons in the *AI* of global efficiency (*E_glob_*) and local efficiency (*E_loc_*). (B) Developmental comparisons in global efficiency and local efficiency within each hemisphere.

To further understand the contribution of lateralization from the developmental view of left and right hemispheric networks, we assessed the group × age interaction effects on the network efficiencies for each hemisphere. We found that the left hemisphere exhibited significant interaction effects on both global efficiency [t = 3.05, p = 0.028] and local efficiency [t = 3.17, p = 0.002], whereas no significant results were found in the right hemisphere [*E_glob_*: t = 1.49, p = 0.14; *E_loc_*: t = 1.81, p = 0.07]. The post hoc analysis further revealed that the TD infants showed significant developmental increases in both global and local efficiencies [*E_glob_*: r = 0.35, p = 0.006; *E_loc_*: r = 0.34, p = 0.008] for the left hemispheres, whereas the infants with congenital SNHL did not exhibit any significant age-related changes in these two efficiency measures [*E_glob_*: r = -0.18, p = 0.18; *E_loc_*: r = -0.24, p = 0.081] (Figure 3B). For the right-hemispheric network, these two groups of infants did not exhibit significant developmental changes in global efficiency [TD: r = 0.17, p = 0.19; SNHL: r = -0.11, p = 0.41] or local efficiency [TD: r = 0.20, p = 0.12; SNHL: r = -0.14, p = 0.32] (Figure 3B). These results demonstrate that the delayed development of network global and local efficiencies in the left hemisphere ultimately led to a developmental deficiency of leftward asymmetry in infants with congenital SNHL.

### 2.4 Hemispheric asymmetry in regional nodal efficiency for each group

The hemispheric asymmetry in nodal efficiency was examined after controlling the age factor. The nodes that showed significant between-hemisphere differences are shown in Figure 4. For the TD infants, leftward asymmetric nodes were mainly located in the frontal and temporal lobes, and one rightward asymmetric node was also found and located in the parietal region. For SNHL infants, leftward asymmetric nodes were primarily located in the frontal lobe, and one rightward asymmetric node was located in the temporoparietal junction region (Figure 4).

**Figure 4.**
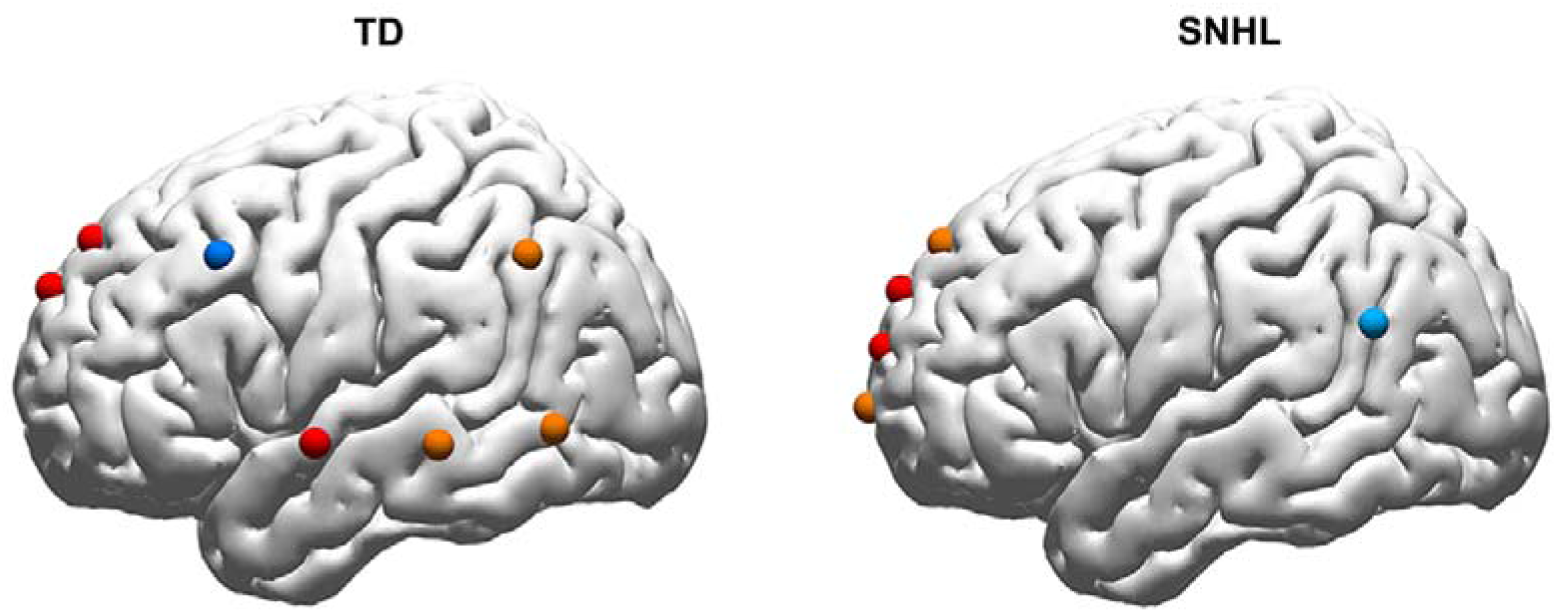
Within-group hemispheric asymmetry in regional nodal efficiency. Red nodes: left > right, significant after correction. Orange nodes: left > right, significant before correction. Dark blue: left < right, significant after correction. Light blue: left < right, significant before correction. p < 0.05 was considered statistically significant.

### 2.5 Developmental differences of hemispheric asymmetry in regional nodal efficiency across groups

A significant group × age interaction effect on the mean *AI* of nodal efficiency was observed in the frontal cortex [t = 2.73, p = 0.007], while other regions did not exhibit such an interaction [temporal: t = 0.54, p=0.59; parietal: t = 0.75, p = 0.46; occipital: t = 0.21, p = 0.83]. Post hoc analysis further indicated that TD infants exhibited a significant developmental increase in the leftward asymmetry of nodal efficiency in the frontal regions [r = 0.35, p = 0.007], whereas such a developmental change was not observed in infants with congenital SNHL [r = -0.11, p = 0.41] (Figure 5A).

**Figure 5.**
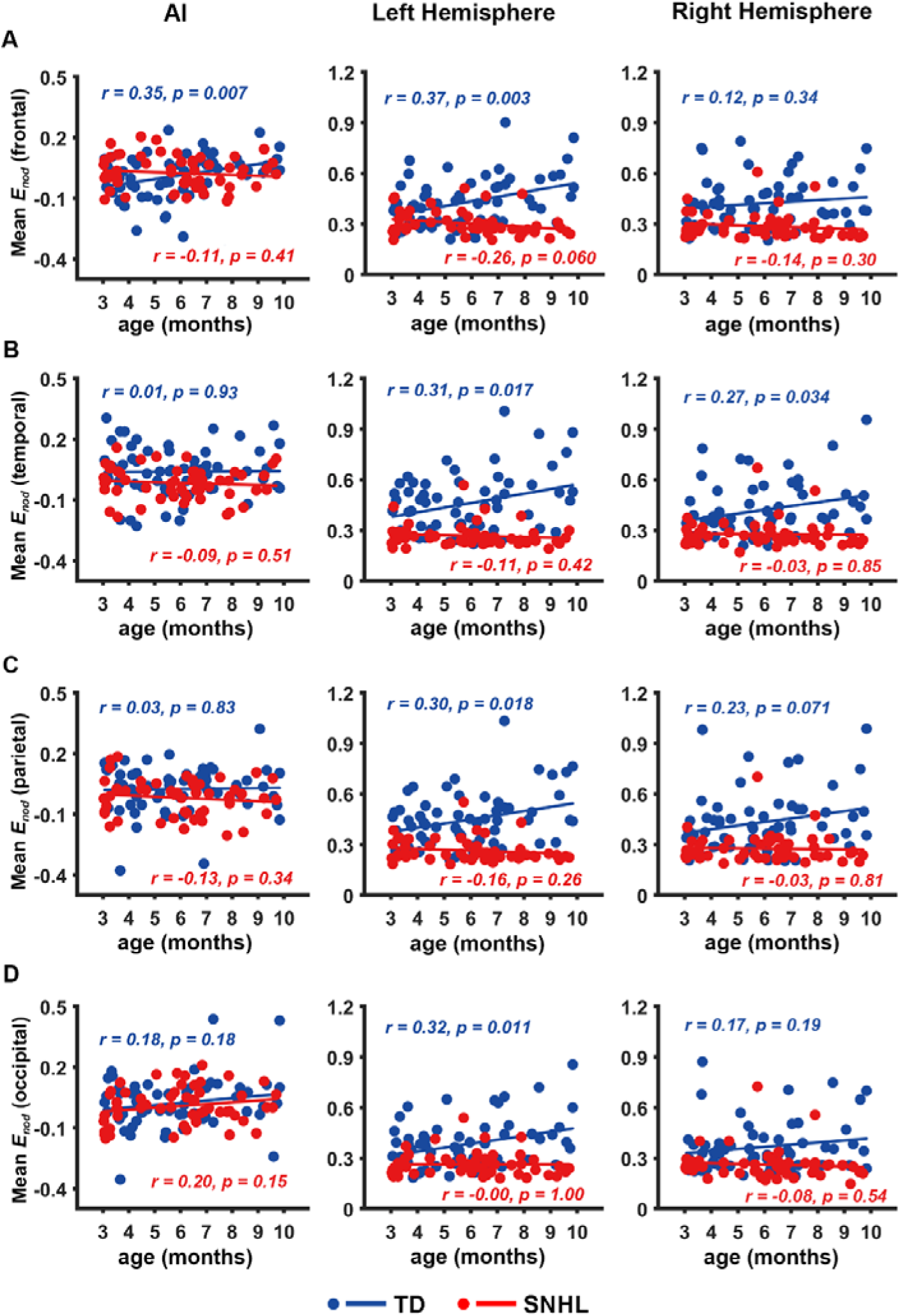
Developmental differences in regional nodal efficiency for TD and congenital SNHL infants. (A-D) Developmental comparisons in the *AI* of nodal efficiency and the mean nodal efficiency within frontal regions of each hemisphere. (A) for the nodes located in frontal regions, (B) for the nodes located in temporal regions, (C) for the nodes located in parietal regions, and (D) for the nodes located in occipital regions.

Furthermore, we investigated the group × age interaction effect on the mean nodal efficiencies of the frontal regions for each hemisphere. A significant interaction effect on the mean nodal efficiencies was observed in the frontal [t = 3.49, p = 0.001], temporal [t = 2.55, p = 0.012], parietal [t = 2.61, p = 0.010], and occipital regions [t = 2.24, p = 0.027] of the left hemisphere. No significant interaction effect on the mean nodal efficiencies was found in the right-hemisphere frontal [t = 1.27, p = 0.21], temporal [t = 1.95, p = 0.054], parietal [t = 1.72, p = 0.088], and occipital [t = 1.40, p = 0.16]. Post hoc analysis revealed that TD infants showed significant developmental increases in the mean nodal efficiency in the left-hemisphere frontal [r = 0.37, p= 0.003] (Figure 5A), parietal [r = 0.31, p = 0.017] (Figure 5B), temporal [r = 0.30, p = 0.018] (Figure 5C) and occipital [r = 0.32, p = 0.011] regions (Figure 5D), but the infants with congenital SNHL did not show significant developmental changes in these left-hemisphere regions [frontal: r = -0.26, p = 0.060; temporal: r = -0.11, p = 0.42; parietal: r = -0.16, p = 0.26] (Figure 5). For the right-hemispheric network, a significant developmental increase in nodal efficiency was observed in the temporal cortex [r = 0.27, p = 0.034] (Figure 5B), and a trend of developmental increase was found in the parietal cortex [r = 0.23, p = 0.071] (Figure 5C) for the TD group, but no significant developmental changes in nodal efficiency were observed in these regions for either the TD group [frontal: r = 0.12, p = 0.34; occipital: r = 0.17, p = 0.19] or the SNHL group [frontal: r = -0.14, p = 0.30; temporal: r = -0.03, p = 0.85; parietal: r = -0.03, p = 0.81; occipital: r = -0.08, p =0.54] (Figure 5).

### 2.6 Association between hemispheric asymmetry and ABR in congenital SNHL infants

The hemispheric asymmetry of global and local network efficiencies did not show any significant correlation with the ABR scores [*AI* of *E_glob_*: r = -0.12, p = 0.37; *AI* of *E_loc_*: r = -0.08, p = 0.54]. After further controlling for age, nonsignificant correlations remained [*AI* of *E_glob_*: r = -0.12, p = 0.38; *AI* of *E_loc_*: r = -0.08, p = 0.57]. These findings indicate that the degree of hearing loss was not necessarily associated with alteration of the degree of network asymmetry during early infancy.

## 3 Discussion

The present study investigates the development of functional network asymmetries in 3- to 9-month infants with congenital SNHL and compares these results to those of age-matched TD controls. These findings show that SNHL infants exhibit lack or delayed development in cerebral asymmetry of functional network efficiencies (e.g., global, local and regional nodal efficiencies) compared to that in TD infants, showing a significant age-related increase in leftward asymmetry. Intriguingly, no significant correlation was observed between the degree of hearing loss and the asymmetry indices of network efficiencies. These results and their implications are discussed in turn.

### 3.1 Economic small-world characteristics of the cerebral network in SNHL infants

Small-world characteristics of the human brain show that the systematic topological network of the cerebral cortex reflects an architecture that optimizes the balance of external and internal functions, indicating that the brain has efficient organization characteristics presumably evolved to support different tasks (Humphries and Gurney 2008; Liu et al. 2020). Earlier findings suggested that this functional brain network organization property exists in brain networks in early life (Huang et al. 2015) and mature over the course of brain development (Cai et al. 2018; Hu et al. 2020). Moreover, previous findings consistently show that the cerebral networks in patients with sensory disorders, such as major depressive disorder (MDD), subjective cognitive decline (SCD), and epilepsy, have small-world properties (Jiang et al. 2019; Lin et al. 2020; Xue et al. 2021). The present study found that both TD and congenital SNHL infants exhibited a higher local efficiency but numerically approximate global efficiency compared with their matched random networks, which is a typical characteristic of small-worldness. The findings of the current study are consistent with previous demonstrations that both the SNHL group and NH group showed small-word organization, with no significant differences between them (Cui et al. 2022). These results demonstrate that the capability of both local specialization and global integration within cerebral networks has emerged and been maintained in TD infants and has not yet been disrupted in the youngest infants with congenital SNHL. Our study suggests that short-term hearing loss does not affect the emergence of small-world characteristics in early infancy, meaning that insufficient auditory exposure has not yet led to disruptive alterations in brain structure and function in infancy.

### 3.2 Aberrant developmental trajectory of hemispheric network asymmetry in SNHL infants

Numerous lines of evidence show that hemispheric network asymmetry emerges in infancy (Dehaene-Lambertz 2002; Taga et al. 2018; Adibpour et al. 2020; Liu et al. 2022) and continues to develop from childhood to adolescence (Szaflarski et al. 2006; Brauer et al. 2008). In line with this, age-dependent abnormalities in brain asymmetry have been reported in adolescents and adults with hearing loss. For example, Allen and colleagues (Allen et al. 2013) reported that congenital hearing loss influenced brain functions, inducing structural changes in extensive cortical regions in adults who used sign language. Twomey and colleagues (Twomey et al. 2017) found that the effect of hearing loss was more task-dependent in the left than in the right superior temporal cortices. However, this outcome was contrary to Payne et al., who found that school-age children with congenital hearing loss aged from 5 to 8 years still showed left asymmetry during language production (Payne et al. 2019). This inconsistency could be explained by a participant’s sign or oral language ability. The infants included in our study were too young to form complete language and comprehension skills. Thus, our results better reflect the effects of congenital hearing loss in the early stages of brain development.

The results of the present study show that TD infants exhibited increased leftward network efficiency asymmetry with age, while SNHL infants aged 3-9 months old did not. These findings are consistent with a previous study showing that both congenitally deaf individuals and those with early acquired deafness (before 3 years old) showed atypical, anomalous cerebral representation (Marcotte and Morere 1990). Collectively, these studies suggest the presence of a critical developmental period for cerebral asymmetry during which adequate auditory exposure is needed to promote left hemispheric superiority for brain organization.

At the regional level, SNHL infants exhibited aberrant development of hemispheric network asymmetry in frontal regions. Several previous resting-state fNIRS reports have elucidated that large-scale brain networks of the frontal region played essential roles in TD infants (Molavi et al. 2014; Cai et al. 2018) and brain regions with significant leftward asymmetry in nodal efficiency in frontal, parietal–occipital junction and occipital regions in TD children and adults (Cai et al. 2019).

Results from the present study suggest that the left hemispheric dominance of small-world network topology was significantly disrupted in the functional cortical networks of infants with congenital SNHL. The disrupted leftward asymmetry in SNHL infants may be explained by the fact that important functional brain network organization properties in the primary visual, auditory, and sensorimotor areas matured around the birth (Bortfeld et al. 2009; Kinsbourne 2009; Paquette et al. 2015). These functional areas manifest a relatively high metabolic rate and are more susceptible to external environmental factors in infants. Meanwhile, the human brain adapts organizationally to environments, physiologic changes, and experiences. Auditory input is detected by cochlear hair cells and transmitted from the cochlear nucleus via the midbrain to the auditory area of the cerebral cortex, which belongs to brain organization. If auditory input is deprived in early life, irreversible morphological changes occur in the human brain (Li et al. 2013). Our results suggest that an age-related increase in leftward asymmetry might underlie the development of a range of brain functions, particularly cognitive and language functions, during early life. Correspondingly, early congenital SNHL appears to inhibit the emergence of these typical asymmetry.

### 3.3 Disrupted development of network efficiency properties in the left hemisphere in SNHL infants

Our results suggest an upward trend in the global and local efficiency of the left hemisphere in TD infants with age, while this fundamental characteristic was impaired in congenital SNHL infants. We observed that the left hemisphere had strengthened information integration and stable transmission function in TD infants with growth, while such brain information processing was disrupted in SNHL infants. Infants with SNHL experience difficulty processing auditory information, which may contribute to the impaired trend in the efficiency of the left hemisphere. This may further result in the difficulties with language and cognitive development observed in these infants’ long term, and may warrant further investigation and intervention to promote optimal brain development.

There are parallels with stronger effects on the left side in noise-induced hearing loss, as described in the literature (Schmidt et al. 2008). Furthermore, we found that the mean *AI* of nodal efficiency and the mean nodal efficiency of the left hemisphere in the frontal lobe improved with age in TD infants at the nodal level. Conversely, congenital SNHL infants exhibited symmetry in the frontal cortex. This result indicated that the maturation of general functions in the left hemisphere was more important than that in the right hemisphere with age.

Previous studies have shown that the left hemisphere of the human brain is dominant in the language processing (Ojemann et al. 1989; Phetsamone et al. 2015) and typically develops faster than the right hemisphere, especially in language-related regions in early life (Qiu et al. 2015; Reynolds et al. 2019). One reason for this difference in development is that language and speech centers tend to develop in this part of brain. This phenomenon is thought to be driven by various factors, such as genetics, how the brain processes information, and exposure to auditory stimuli during cortical developmental periods. Our study provides new evidence that adequate auditory exposure is vital for brain development.

### 3.4 Lack of auditory exposure in early life affects brain development

Numerous studies have shown that the human brain is considered environment dependent. The development of the auditory cortex extensively depends on hearing experience (Kral & Sharma, 2012; Schaafsma et al., 2009). Insufficient stimulation of the auditory cortex could affect multisensory systems such as the function of audition, language, and vision. A report on individuals with congenital hearing loss who used sign language also showed evidence of structural cortical changes in extensive brain regions, indicating that specific environments influenced brain functions (Allen et al. 2013). The current study provided new evidence for this debate. Our results showed that there was no significant correlation between the degree of hearing loss and *AI* in congenital SNHL infants, which indicated that even mild hearing loss could disrupt the development of cerebral asymmetry. This finding emphasized the importance of ensuring that infants are exposed to sufficient auditory exposure through early life. A previous study suggested that the hemispheric asymmetry pattern for auditory information processing was associated with the severity of hearing impairment in adults (Zhu et al. 2020). Subcortical disease studies have suggested that interhemispheric asymmetry is related to the degree of clinical impairment in adults, such as ischemic stroke (Fanciullacci et al. 2017). However, there are few reports on the correlation between the asymmetry of the brain network and the degree of hearing loss during infancy. The observed lack of correlation between the severity of hearing loss and hemispheric asymmetry might be explained by the highly plastic critical functional brain network organization in early childhood and environmental factors being critical to the development of the cerebral cortex. If infants receive insufficient auditory stimulation, even with mild hearing loss, the brain network will develop abnormally, especially in the dominant hemisphere.

It is important to note that the impact of SNHL on hemispheric network asymmetry and language development can vary greatly depending on the severity and timing of hearing loss, as well as the availability of early interventions such as hearing aids or cochlear implants. Early identification and intervention can help to support the development of typical hemispheric network asymmetry and minimize the impact of SNHL on language and communication development in infants. The recovery of functional network asymmetry in infants with early intervention needs to be investigated to consolidate the primary findings.

Our results also revealed that age-related development in the *AI* of the global and local network efficiencies was not observed in congenital SNHL infants, which suggested that early SNHL caused atypical changes in cortical function. This finding is consistent with previous studies on older children with bilateral cochlear implants (Cis) (Sevy et al. 2010; Gordon et al. 2013). Several reports have shown that early intervention for hearing loss, such as hearing aids (HAs) and CIs, induced brain reorganization and plasticity (Anderson et al. 2017). The existing body of research suggests that left hemisphere dominance for language was found in children with CI and controls (Chilosi et al., 2014). The asymmetry of cortical functional areas represents the best prerequisite for learning speech, language, and other skills. Early identification of patients who exhibit atypical development of cortical function is critical for early intervention and preventing delays in linguistic and psychosocial development.

We demonstrated the impact of short-term insufficient auditory exposure on the hemispheric brain network, which is a cross-sectional study. However, the hemispheric asymmetry of the brain network may change differently with the input of auditory stimuli. Longitudinal studies are needed to validate our results. In the future, the debate on the causes of brain asymmetry is intriguing and could be usefully explored in further research.

The mean age of congenital SNHL infants was 5.83 months in our study, and their brain cortical functions had already been impacted. Hence, these congenital SNHL infants should engage in early intervention as soon as possible. Interestingly, this is consistent with the recommended time of hearing impairment intervention per guidelines, which is 6 months. Therefore, fNIRS may provide a new auxiliary clinical tool for personalized early intervention. Furthermore, the present study showed that fNIRS is sensitive to monitoring cortical function, which has also been a popular addition to a limited choice of neuroimaging modalities suitable for CI recipients. Therefore, fNIRS has the potential to provide an additional clinical assessment of the changes in cortical function induced by auditory stimuli. Anderson et al. predicted CI outcomes in deaf adults by comparing pre-and postoperative brain activation measures using fNIRS (Anderson et al. 2017). A review also proposed using fNIRS to supplement the current clinical practice of CI programming and speech and language therapy (Saliba et al. 2016). Our present results provide potential directions for future fNIRS applications in assessing and evaluating the early personalized intervention.

Overall, a lack of adequate auditory exposure may alter the information transfer patterns of functional networks, and if this condition persists for a long time, the brain will adjust to adapt and compensate for the lack of auditory exposure to maintain topological organization. These results highlight the importance of early identification and intervention for infants with SNHL, as well as the need for further research to better understand the impact of SNHL on cerebral asymmetry and functional network efficiency. By improving our understanding of the effects of SNHL on brain development, we may be able to develop new strategies to support language and communication development in this population. Further research is needed to better understand the complex relationships between SNHL individuals, hemispheric network asymmetry, and language development.

## 4 Conclusion

In summary, the current study demonstrated that hemispheric asymmetry formed in early life and the initial lack of effective auditory exposure affected the typical development of brain network in SNHL infants. Although the infants with congenital SNHL exhibited a balance between information segregation and integration within two hemispheres, consistent with the TD controls, the development of hemispheric asymmetry in network efficiency measures was disrupted. These findings indicated that insufficient auditory cortex stimulation in early life could affect the development of functional network architecture in specialized cortical regions. Our current study filled a gap in research on functional topological asymmetry between hemispheres in infants with early congenital SNHL. Meanwhile, our study provided a basis for follow-up research on the cortical function of infants with congenital SNHL and some theoretical bases for early auditory personalized intervention.

## 5 Materials and Methods

### 5.1 Participants

Fifty-five congenital SNHL infants aged 3 to 9 months and 60 age- and gender-matched TD infants participated in this study. Table 1 provides a summary overview of the participant’s demographic characteristics. Congenital SNHL infants were all recruited from Beijing Children’s Hospital, Capital Medical University. Of the TD infants, 20 were recruited from Beijing Normal University, while the remaining 40 were recruited from Beijing Children’s Hospital, Capital Medical University. All participants were recruited following the same procedure. Informed consent was obtained from infants’ parents, with experimental protocols approved by the Institutional Review Board of Beijing Children’s Hospital, Capital Medical University and the Institutional Review Board of State Key Laboratory of Cognitive Neuroscience and Learning, Beijing Normal University.

**Table 1.**
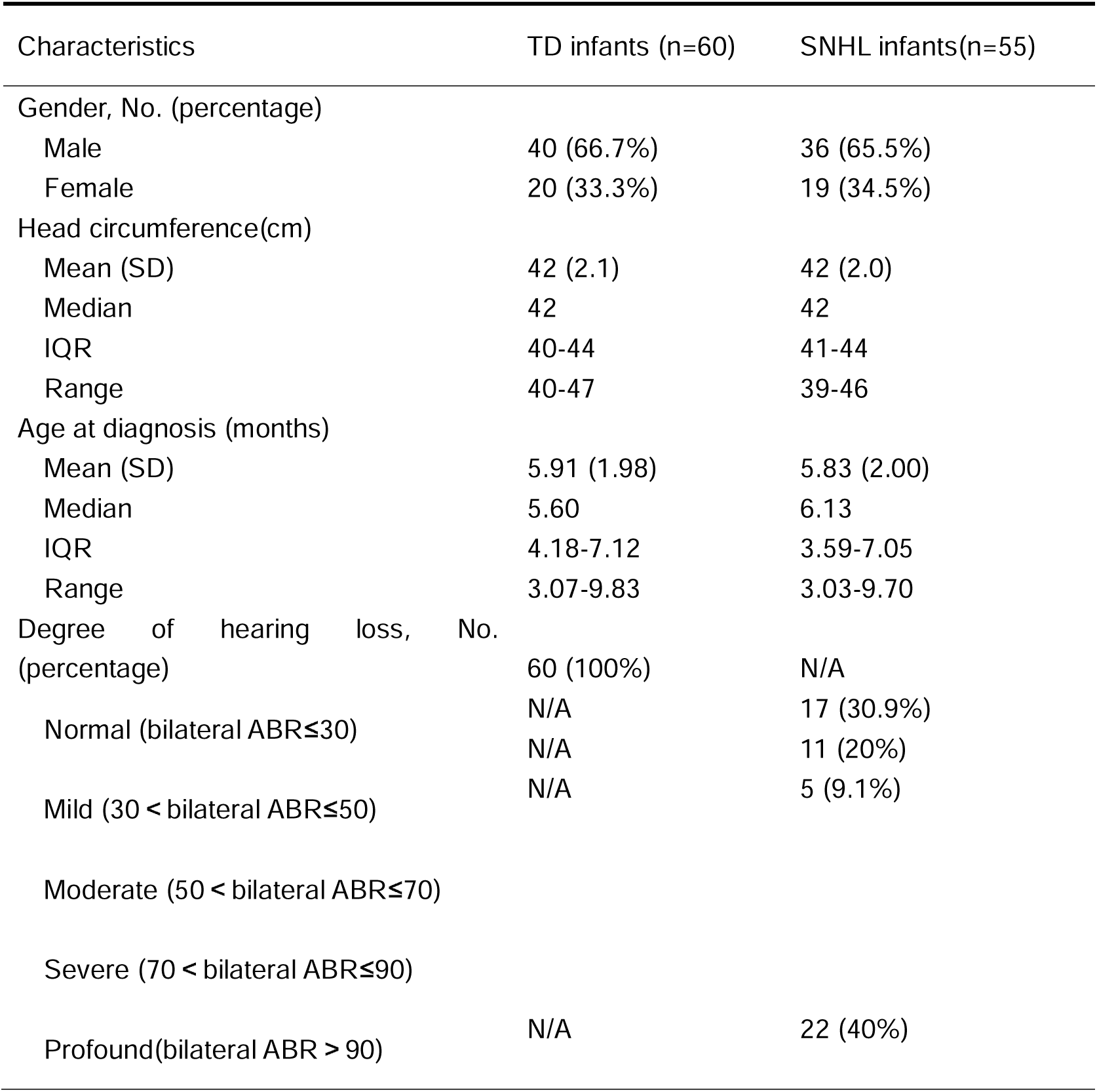
Demographic characteristics of all participants.

All infants received a full audiological diagnosis guided by the cross-check principle. The diagnosis test battery included auditory brainstem response (ABR), auditory steady-state response (ASSR), distortion product otoacoustic emission (DPOAE), and acoustic impedance. Inclusion criteria for the congenital SNHL infant group were as follows: (1) bilateral thresholds of ABR Wave V**>**30 dB nHL; (2) the average threshold of ASSR**>**30 dB nHL at 0.5, 1, 2, 4 kHz; (3) bilateral 1-kHz tympanograms were single-peaked and 0.226-kHz tympanograms were Type A; (4) bilateral DPOAE were absent at least four frequencies. Inclusion criteria for the TD infants were: (1) bilateral thresholds of ABR Wave V≤30 dB nHL; (2) an average threshold of ASSR≤30 dB nHL at 0.5, 1, 2, 4 kHz; (3) bilateral 1-kHz tympanograms were single-peaked and 0.226-kHz tympanograms were Type A; (4) bilateral DPOAE were present in at least four frequencies. Infants had no other known cognitive or neurological impairments, developmental delay, or active external or middle ear infections.

### 5.2 NIRS data recording

A continuous wave near-infrared optical imaging instrument (NirSmart, Hui Chuang, China) with a sampling rate of 10 Hz was used to measure hemodynamic changes in the cerebral cortex of the infants. Oxygenated hemoglobin (HbO) and deoxygenated hemoglobin (HbR) have different absorption spectra in the near-infrared (NIR) wavelength region. The NIRS instrument exploits these optical properties of hemoglobin by using two wavelengths of light, 760 nm and 850 nm, which are differentially absorbed by the two chromophores. The probe array was composed of 24 sources and 16 detectors interlacing at the nearest distance of 3 cm, generating a total of 64 measurement channels covering the frontal, temporal, parietal, and occipital areas of each hemisphere (Figure 6A).

**Figure 6.**
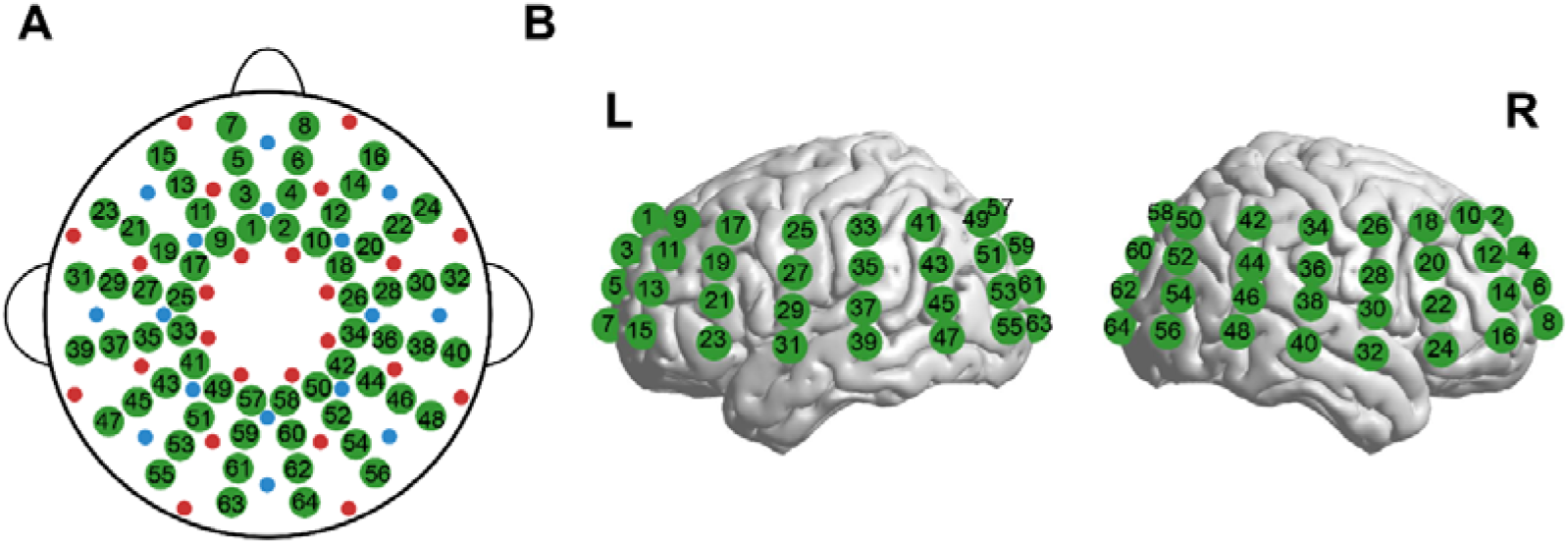
fNIRS probe arrangement and MRI neuroanatomical coregistration. (A) The arrangement of the 64 measurement channels. The red and blue dots represent the sources and detectors, respectively. The green dots with digits represent the positions of the measurement channels. (B) MRI coregistration was conducted by an infant aged 6 months old wearing probe arrays with vitamin E capsules in MRI. Green dots with digits represent the measurement channels covering each hemisphere’s frontal, temporal, parietal and occipital areas.

The experiment was conducted in a dimly lit, quiet room. The probe array was symmetrically positioned on the infants’ scalp following the 10–20 system, with reference to the external auditory canals, inion, and vertex as landmarks. Infants were held by their mothers during data collection, and mothers were instructed to remain silent and minimize movements during data acquisition to prevent displacement of the probes. For each infant, fNIRS data were collected for approximately eleven minutes while the infant was asleep.

### 5.3 MRI coregistration

To validate the position of the probes on the scalp, a structural MR image (MRI) was acquired from a 6-month-old infant wearing probe arrays with vitamin E capsules. Vitamin E locations from these MRI scans were used as landmarks for coregistration, a common way to localize fNIRS probes relative to the brain (Kovelman et al. 2008; Sasai et al. 2012). The scan was performed at the Imaging Centre in the Beijing Children’s Hospital using a 3T Siemens Tim Trio MRI scanner. T1-weighted structural images were acquired using a magnetization-prepared rapid gradient echo (MPRAGE) sequence. The scanning parameters were as follows: slices=152, TR=6.67 ms, TE=3.02 ms, FOV=202×202 mm, voxel size=1 mm×1 mm×1 mm, flip angle=8°, and slice orientation=sagittal. The sMRI data were normalized into Montreal Neurological Institute (MNI) space using a 6-month-old infant’s brain template, and location coordinates were projected on the brain surface using NIRS-SPM software (http://bispl.weebly.com/nirs-spm.html) (Ye et al. 2009). The channels covered the frontal, temporal, parietal, and occipital areas of each hemisphere according to the automated anatomical labeling (AAL) template (Tzourio-Mazoyer et al. 2002) [Figure 1(B)]. Detailed cortical positions corresponding to each NIRS channel are shown in Table S1 in Supplementary Materials.

### 5.4 Data preprocessing

#### Quality control

Similar to our previous study (Liu et al. 2022), we also used the in-house FC-NIRS package (Xu et al. 2015) (https://www.nitrc.org/projects/fcnirs) to preprocess fNIRS data in this study. First, we examined the signal quality for each participant using three different approaches: signal-to-noise ratio (SNR), between-channel signal correlation, and frequency spectrum analysis. Generally, a measurement signal with low SNR (< 2) was considered to be a “bad channel” reflecting poor contact between the optodes and scalp. Such bad channels are usually characterized by a low correlation between this signal and other signals, as assessed using signal correlation analysis. They also generally lack a cardiac component in the frequency spectrum analysis. We implemented these criteria as the basis for the removal of bad channels with poor signal quality. The number of channels removed ranged from 0 to 4 for SNHL infants and from 0 to 4 for TD infants.

#### Reducing motion artifacts and global interference

Next, we detected and corrected motion-induced artifacts using the spline interpolation method (Scholkmann et al. 2010). This approach calculates the moving standard deviation (MSD) within sliding windows of 2-second lengths. MSD values larger than 5 standard deviations from the mean were considered artifacts. The time series representing motion artifacts were further modeled via a cubic spline interpolation and subtracted from the original time series. The resulting signal was considered to be free of motion artifacts. Subsequently, we extracted a relatively stable time series from the processed data for each participant. Furthermore, we filtered out the first principal component through a principal component analysis (PCA) algorithm, in order to reduce the effect of global noise, which is generally associated with systemic fluctuations (e.g., arterial pulse, respiration, and cardiac pulsation) (Franceschini et al. 2006; Abdalmalak et al. 2022).

#### Calculating low-frequency hemoglobin signals

Finally, we bandpass-filtered the data between 0.009 and 0.08 Hz to remove low- and high-frequency noise, such as cardiac noise and slow drift (Liu et al. 2022), and calculated the relative changes in the concentrations of oxyhemoglobin (HbO) and deoxyhemoglobin (HbR) based on the modified Beer-Lambert law (Kocsis et al. 2006). We used HbR signal to characterize topological characteristics of functional networks because HbR signal is the physiological basis of the blood oxygen level-dependent fMRI signal (Ogawa et al. 1993; Buxton et al. 1998) and has been shown to have higher test-retest reliability in an fNIRS-based brain network study (Niu et al. 2013).

### 5.5 Hemispheric network construction

In the context of fNIRS, network nodes were defined as measurement channels, and edges were defined as functional connectivity between nodes as estimated by a Pearson correlation of the time series. To investigate the network asymmetry of infants’ brains, we constructed the left- and right-hemispheric brain networks for each participant based on the within-hemispheric nodes. Given the ambiguous biological basis for negative correlations in functional connectivity analyses, we set all negative correlations to zero and restricted our analysis to positive correlations. To improve the normality distribution of the correlations in the matrix, we converted correlation coefficients into z values via Fisher’s r-to-z transformation. As in our previous studies (Cai et al. 2019; Liu et al. 2022), to ensure that the networks have the same number of edges for each infant, we thresholded each correlation matrix into a weighted matrix with a fixed sparsity value, defined as the total number of edges in a network divided by the maximum possible number of edges. We chose sparsity (S) over a range of 0.2-0.5 with an interval of 0.01 for theoretical graph analysis of the hemispheric brain network.

### 5.6 Topological network measures

#### 5.6.1 Network efficiency properties

In graph theory, topological network efficiencies are often used to demonstrate efficient information processing within complex networks (Latora and Marchiori 2001, 2003). Meanwhile, these efficiency measures are conceptually preferable for characterizing brain network topology (Achard and Bullmore 2007). We, therefore, focused on efficiency-related network properties in our investigation of the brain networks in both SNHL and TD infant groups. Specifically, global network efficiency, local network efficiency, and nodal efficiency were computed within each hemisphere for each participant. The definitions for these parameters are described below.

##### Network global efficiency

The global efficiency *E_glob_* of network *G* is the average inverse shortest path length of all node pairs (Latora and Marchiori 2001) and was calculated as follows:

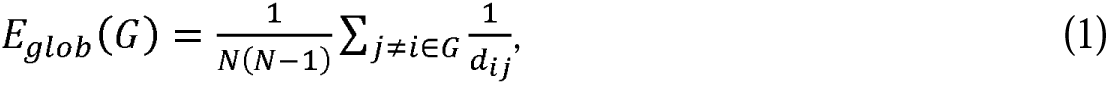

where *d_ij_* is the shortest path length between node *i* and node *j*, and *N* is the number of nodes in the network.

##### Network local efficiency

Network local efficiency is defined as the efficiency of the local subgroup of node *i*, which consists of only the direct neighbors of node *i*. (Latora and Marchiori 2001; Achard and Bullmore 2007). The local efficiency of network *G* is calculated as follows:

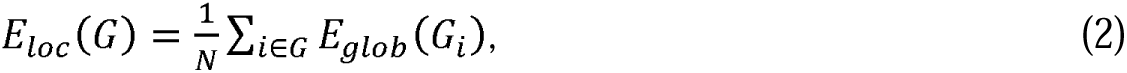

where *E_glob_* (*G_i_*) is the global efficiency of *G_i_*, the subgraph of the neighbors of node *i*.

##### Nodal efficiency

The efficiency of the node was defined as the harmonic mean of the shortest path length between this node and all its neighbors(Latora and Marchiori 2001) and is measured as follows:

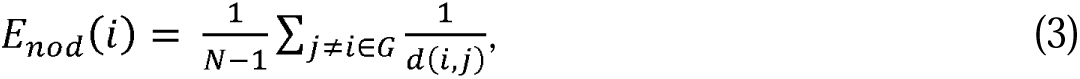

where *d*(*i, j*) is the shortest path length between node *i* and node *j*.

These network properties were calculated using our in-house FC-NIRS package (Xu et al. 2015, https://www.nitrc.org/projects/fcnirs), and the results reported in this study were based on sparsity of 0.45.

#### 5.6.2 Small-world efficiency analysis

Small-world properties are fundamental characteristics of human brain networks, describing the capability of efficient global information integration and local information processing within the brain. Small-world network properties were originally proposed by Wattz and Strogatz (1998). Here, we employed global network efficiency and local network efficiency to quantify the small-world behavior of cortical networks in SNHL and TD infants, respectively. These network efficiency measures were compared separately to the corresponding indices derived from 1000 comparable random null networks. The random networks were generated by preserving the same numbers of nodes and edges and the same degree distribution as the brain network. Typically, a small-world network should meet the following criteria: global efficiency in a real brain network is numerically approximate to that in a random network, and local efficiency in the brain network is larger than that in a random network. To investigate the small-world topology in SNHL infants and TD infants, we computed normalized *E_loc_* and normalized *E_glob_*, defined as the ratio of network efficiencies from hemispheric networks to that from their matched random networks across a wide sparsity range (0.2∼0.5).

#### 5.6.3 Efficiency asymmetry calculation

To quantify the degree of asymmetry in efficiency measures between hemispheres, we calculated a commonly used asymmetry index (*AI*), which was defined as follows:

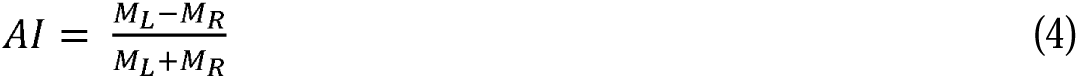

For network efficiencies, *M_L_* and *M_R_*represent the global or local efficiency of the entire left and right hemispheric network, respectively. For nodal efficiencies, *M_L_* and *M_R_* represent the mean nodal efficiency within different brain regions (frontal, temporal, parietal, and occipital regions) from the left and right hemispheric networks, respectively. The *AI* values range from -1 to 1, with a positive value of *AI* representing leftward asymmetry and a negative value of *AI* representing rightward asymmetry.

### 5.7 Statistical analysis

To investigate the developmental difference in global network efficiency, local network efficiency and nodal efficiency between the SNHL and TD infant groups, we performed a general linear model (GLM) with “age”, “group”, and “group × age” as predictor variables on each efficiency measure. A significant group × age interaction suggests a difference in brain network efficiency between two groups.

To evaluate the hemispheric difference in global efficiency, local efficiency and nodal efficiency between the left and right hemispheres within the TD and congenital SNHL infant groups, we adopted a mixed-effect linear model for each metric within each group, with hemisphere taken as a repeated measure (i.e., left and right as repeated), efficiency parameters taken as prediction factors, and gender and age included as covariates. For nodal efficiencies (32 nodes in each hemisphere), the false discovery rate (FDR) procedure was applied to correct for multiple comparisons. Of note, p < 0.05 after the correction was considered significant.

Next, we assessed age-related effects on the *AI* of the network’s global efficiency, local efficiency, and nodal efficiencies. A general linear model (GLM) was used with “age”, “group”, and “group × age” as predictor variables for each *AI* measure. A significant group × age interaction on the *AI* suggested a difference in brain asymmetry between two groups. To ascertain how the two groups differed in the developing trajectory of *AI*, we evaluated correlations between the *AI* and age within each group. We also used a general linear model (GLM) with “age”, “group”, and “group × age” as predictor variables on the topological parameters for each hemisphere to explore how the two hemispheres differ in the development of network efficiencies. After that, we evaluated the correlation between the efficiency within each hemisphere and age to interpret the effect of each hemisphere on development.

## Acknowledgments

This study is supported by the National Natural Science Foundation of China (81761148026 and 81571755) and the Open Research Fund awarded to the State Key Laboratory of Cognitive Neuroscience and Learning at Beijing Normal University.

